# A single-cell atlas of the single versus multiple parous Hu Sheep ovary

**DOI:** 10.1101/2023.05.22.541677

**Authors:** Ting Ge, Yifan Wen, Bo Li, Xiaoyu Huang, Shaohua Jiang, Enping Zhang

**Affiliations:** College of Animal Science and Technology, Northwest A&F University, Xianyang 712100 Shaanxi, China

**Author notes:** Corresponding author. EM.

**Keywords:** Hu Sheep, Single-cell RNA sequencing, Lambing number, Ovarian somatic cells, Granulosa cells

## Abstract

In the modern sheep production system, the reproductive performance of ewes determines the economic profitability of farming. The mechanism of difference in litter size is important for the selection and breeding of high-fecundity ewes. Hu sheep is a high-quality sheep breed with high fecundity in China and is ideal for investigating high reproductive traits. In the current study, the sheep with lambing number ≥3 in three consecutive lambing records were assigned to the HLS group, and lambing number = 1 as the LLS group selected from the same farm with three consecutive lambing. Three randomly picked ewes were slaughtered within 12 h of estrus, and unilateral ovarian tissue was collected and analyzed by single-cell RNA sequencing in each group. A total of five types of somatic cells were identified, and corresponding expression profiles were mapped in the ovaries of the Hu sheep. Additionally, the results of the difference in ovary somatic cell expression profiles between HLS and LLS present that the differences between multiples vs. singleton Hu sheep were mainly clustered in the GCs. In addition, 4 granulosa cell subtypes were identified. GeneSwitches results revealed the opening of *JPH1* expression and the closure of *LOC101112291*, which leads to different evolutionary directions of the granular cells. The expression levels of *FTH1* and *FTL* in GCs of Hu sheep in the HLS group were significantly higher, which inhibited necroptosis and ferroptosis of mural– GCs from decreasing follicular atresia. This study constructed the cellular atlas of the ovary and revealed related biological characteristics at the cellular molecular level. It provides a theoretical basis for the mechanisms underlying the differences in ovulation numbers, which contributes to breeding high-fertility sheep and molecular genetics-based selection.

## 1. Introduction

For a long time, sheep have undergone a strong selection pressure to improve fecundity because high reproduction trait always brings high economic merit. The ovary is a reproductive organ in sheep consisting of follicles at several different developmental stages. The number of lambs produced in sheep is an important indicator of sheep fertility. It is also a complex quantitative trait regulated by genetic, epigenetic, and hormonal factors with a heritability of 0.03-0.10 [1]. The number of lambs produced by ewes is influenced by the number of ovulations, and ovulation can be genetically regulated by a single main effector gene and some micro-effector polygenes [2, 3], such as *BMPRIB* [4], *BMP15* [5], and *GDF9* [6].

The ovary is a heterogeneous organ co-regulated by multiple cells, which determines the complexity of the ovarian function. Follicular development is a highly coordinated process in Hu sheep. Follicle-cyclic recruitment, spatial displacement, follicle atresia, and ovulation are implicated events resulting from the somatic cells’ release of molecular signals. Cells have different functions in the specific biological cycle of the ovary and contribute to the maturation of follicles [7]. Previous studies have taken the ovary as a whole [8, 9]. Few studies have been conducted to investigate the effect of different cells in the ovary on reproductive performance. Therefore, establishing a functional analysis based on different cells of the ovary and their specific physiological roles and molecular mechanisms is important to explore the ovarian function and elucidate the mechanisms of differences in lambing number.

With the development of sequencing technology, single-cell RNA sequencing (scRNA-seq) technology has been proposed to detect the expression profiles of different tissue cells. Thousands of single-cells from a single biopsy can be analyzed by introducing unique molecular identifiers (UMI) in droplet-based protocols, reducing amplification errors and facilitating the detection of small populations of cells whose transcriptional programs are often not detected using bulk RNA sequencing [10]. Many studies have focused on using single-cell transcriptome technology to understand the different cell functions and developmental trajectories of the ovary [11–13]. At distinct developing stages of cells, the corresponding cell markers are different [14–16]. In the current study, the scRNA-seq technique explored the mechanisms of different lambing number differences based on the cellular level.

Hu sheep is a first-class protected local livestock breed in China and a world-renowned multiparous sheep breed. It has early sexual maturity, four seasons of estrus, two or two years of three litters a year, and an average lambing rate of 277.4%, with more than 6 to 8 ewes [17]. Hu sheep are currently farmed on a large scale in China’s mutton sheep production system, and the litter size of ewes impacts economic efficiency. In sheep breeds with high fecundity performance, five main effect genes control ovulation and lambing numbers. However, except for the *FecB* locus in the *BMPR-1B* gene, all other loci are not associated with high fecundity traits in Hu sheep [18]. There is still a gap between the specific gene regulatory networks and lamb number of different mechanisms. In this study, we focused on the critical window period of estrus through 10×Genomics single-cell sequencing technology to uncover the key biological processes of cells in the ovaries of single/multi-lamb Hu sheep to investigate the key genes and pathways associated with the lambing number in the ovaries, which provide new targets for molecular breeding and theoretical basis for further studies.

## 2. Materials and methods

### 2.1 Ethical statement

The protocol of this study was approved by the Animal Care and Use Committee of Northwest A&F University, China (Approval No. DK2021113).

### 2.2 Sheep management

A total of 6 estrus Hu sheep were divided into 2 groups, and litter size from three consecutive parities was used as a grouping basis. High-reproduction group (HLS) had a litter size of ≥ 3, and the low-reproduction group (LLS) had 3 sheep which had a litter size of =1. Sheep weight and production records are shown in Table s1.

Ewe were slaughtered within 12 h of estrus. The ram test was used to determine the estrous status. Venous blood was collected before slaughter for testing blood biochemical and hormone levels. Unilateral ovarian tissue was collected, placed in a protective solution (MACS® Tissue Storage Solution, **Miltenyi Biotec Inc, GER**), stored at 4 ℃, and analyzed by scRNA-seq.

### 2.3 Blood biochemical and hormone determination

Blood biochemical indicators were measured with an automatic biochemical analyzer (BK-280, Biobase, CHN). Blood hormone concentration was tested according to the instructions of the Elisa kit, and the kit information is shown in Table S2.

### 2.4 Single cell RNA sequencing

#### 2.4.1 Sample preparation

The entire unilateral ovary of Hu sheep was cut up, digested with collagenase1 for 30 min and trypsin for 10 min, sieved, centrifuged, and lysed for cell counting. The 10x Genomics Chromium system (10x Genomics, USA) used an 8-channel microfluidic “double cross” crossover system to mix barcode-containing gel beads, a mixture of cells and enzymes, and oil to form GEMs (a water-in-oil microsystem). After forming GEMs, the cells were lysed, and the gel beads automatically dissolved to release many barcode sequences. The oil droplets were ruptured, and complementary DNA (cDNA) products were collected and amplified. Total messenger RNA (mRNA) was reverse transcribed to produce cDNA with 10x barcode and UMI information. A standard sequencing library was constructed, and the 10x library was cyclized and sequenced using the MGISEQ sequencing platform (MGI Tech Co., Ltd, China).

#### 2.4.2 Sequencing

Single-cell RNA-seq libraries were prepared with Chromium Single cell 3’ Reagent v2 (or v3) Kits (10x Genomics, USA) according to the manufacturer’s protocol. Single-cell suspensions were loaded on the Chromium Single Cell Controller Instrument (10×Genomics, USA) to generate single-cell gel beads in emulsions (GEMs). After generating GEMs, reverse transcription reactions were engaged with barcoded full-length cDNA followed by the disruption of emulsions using the recovery agent and cDNA clean-up with DynaBeads Myone Silane Beads (Thermo Fisher Scientific, Waltham, MA, USA). cDNA was amplified by polymerase chain reaction (PCR) with appropriate cycles, depending on the recovery cells. The amplified cDNA was fragmented, end-repaired, A-tailed, index adaptor-ligated, and library amplification. These libraries were sequenced on the Illumina sequencing platform (HiSeq X Ten), and 150 bp paired-end reads were generated.

#### 2.4.3 Data preprocessing

The Cell Ranger software pipeline (version 5.0.0) provided by 10×Genomics was used to demultiplex cellular barcodes, and reads were mapped to the genome and transcriptome using the STAR aligner and down-sample reads as required to generate normalized aggregate data across samples, producing a matrix of gene counts versus cells. The unique molecular identifier (UMI) count matrix was processed using the R package Seurat [19] (version 3.1.1). To remove low-quality cells and multiplet captures, a major concern in microdroplet-based experiments, a criterion was applied to filter out cells with gene numbers less than 200, UMI less than 1000, and log10GenesPerUMI less than 0.7. We discarded low-quality cells where >10% of the counts belonged to mitochondrial genes, and >5% of them belonged to hemoglobin genes. The DoubletFinder package [20] (version 2.0.2) was applied to identify potential doublet. After applying these QC criteria, 41150 single-cells were included in downstream analyses. Library size normalization was performed with the NormalizeData function in Seurat [18] to obtain the normalized count. The global-scaling normalization method “LogNormalize” normalized the gene expression measurements for each cell by the total expression and multiplied by a scaling factor (10,000 by default). The results were log-transformed.

Top variable genes across single-cells were identified using the method described in Macosko et al. [21]. The most variable genes were selected using the FindVariableGenes function (mean.function = FastExpMean, dispersion.function = FastLogVMR) in Seurat [19]. Principal component analysis (PCA) was performed to reduce the dimensionality with the RunPCA function in Seurat [19]. Graph-based clustering was performed to cluster cells according to their gene expression profiles using the FindClusters function in Seurat [19]. Cells were visualized using a 2-dimensional Uniform Manifold Approximation and Projection (UMAP) algorithm with the RunUMAP function in Seurat [19]. The FindAllMarkers function (test.use = presto) was used in Seurat [19] to identify marker genes of each cluster. For a given cluster, FindAllMarkers identified positive markers compared with all other cells.

Differentially expressed genes (DEGs) were identified using the FindMarkers function (test.use = presto) in Seurat. *P* value < 0.05 and |log2foldchange| > 0.58 were set as the threshold for significantly differential expression. Gene Ontology (GO) enrichment and Kyoto Encyclopaedia of Genes and Genomes (KEGG) pathway enrichment analysis of DEGs were performed using R based on the hypergeometric distribution.

#### 2.4.4 Pseudotime analysis

Pseudotime analysis was done with the Monocle2 package [22]. The raw count was converted from the Seurat object into the CellDataSet object with the import CDS function in Monocle. The differentialGeneTest function of the Monocle2 package was used to select ordering genes (qval < 0.01), which were informative in the ordering of cells along the pseudotime trajectory. The dimensional reduction clustering analysis was performed with the reduce dimension function, followed by trajectory inference with the order. The cell function was done using default parameters. Gene expression was plotted with the plot_genes_in_pseudotime function to track changes over pseudo-time.

#### 2.4.5 GeneSwitches analysis

GeneSwitches (v 0.1.0) [23] was used to discover the sequence of gene expression turn-on, and turned-off during cell state transitions at single-cell resolution. Gene expression data were binarized to a 1 (on) or 0 (off) state using the binarize_exp function (fix_cutoff = TRUE, binarize_cutoff = 0.05) from the GeneSwitches package. A mixture model of two Gaussian distributions was fitted to the input gene expression for each gene, which was used to calculate a threshold for binarizing the gene. Genes without a significant “on-off” bimodal distribution were removed, and the binary state of gene expression (on or off) was modeled using the find_switch_logistic_fastglm function (downsample = TRUE). The top 50 best-fit (high McFadden’s Pseudo R^2) genes were plotted along the proposed timeline. Genes turned on with the proposed time were shown above the horizontal axis. Genes that were turned off with the proposed time are shown below the horizontal axis.

### 2.5 Histological observation

#### 2.5.1 Hematoxylin and eosin staining

The ovary samples were fixed in 4% polyformaldehyde, embedded in paraffin, sectioned, and stained with hematoxylin and eosin (H&E) for histologically observing ovarian tissues. SlideViewer 2.5.0 (3DHISTECH Ltd., Hungary) was used for imaging.

#### 2.5.2 Immunofluorescence staining

The sections of paraffin-embedded tissues were stained with Decorin antibody (sc-73896, Santa Cruz, USA), VE-cadherin antibody (sc-9989, Santa Cruz, USA), CD53 antibody (sc-9989, Santa Cruz, USA), FSHR antibody (22665-1-AP, Proteintech, CHN), RGS5 antibody (11590-1-AP, Proteintech, CHN), and DAPI (C0060, Solarbio, CHN) for immunofluorescence. SlideViewer 2.5.0 (3DHISTECH Ltd., Hungary) was used for imaging.

### 2.6 Statistical analysis

The Student’s *t*-test was used for data analysis by SPSS software, version 24.0. (SPSS, Chicago, IL, USA). Statistical significance was set at *p* < 0.05 and extremely significant at *p* < 0.01.

## 3. Results

### 3.1 Blood biochemical and body weight of sheep

12 hours before sampling the ovary, the venous blood was collected to evaluate the physiological status of the sheep. As shown in Table 1, the blood biochemical and body weight of sheep had no difference between the two groups.

**Table 1.** Blood biochemical and body weight of sheep.

| Item | LLS | HLS | *p*-value |
| --- | --- | --- | --- |
| TP, g/L | 70.70±2.61 | 76.03±2.41 | 0.923 |
| GLU, mmol/L | 3.73±0.29 | 3.27±0.17 | 0.339 |
| TC, mmol/L | 1.98±0.27 | 1.57±0.26 | 0.980 |
| HDL-C, mmol/L | 1.41±0.44 | 0.99±0.24 | 0.373 |
| LDL-C, mmol/L | 0.80±0.10 | 0.65±0.09 | 0.953 |
| TG, mmol/L | 0.17±0.04 | 0.20±0.05 | 0.571 |
| E<sub>2</sub>, ng/L | 58.50±6.38 | 58.59±1.49 | 0.060 |
| LH, ng/L | 17.75±2.44 | 13.84±0.57 | 0.210 |
| FSH, IU/L | 32.53±2.98 | 29.93±3.84 | 0.511 |
| T, nmol/L | 7.42±0.19 | 5.86±0.10 | 0.328 |
| Prog, pmol/L | 941.88±46.44 | 1057.59±152.12 | 0.057 |
| Weight, Kg | 42.01±0.31 | 40.16±1.19 | 0.097 |

### 3.2 Clustering and identification of the ovary somatic cells

In this study, ovaries were obtained from 6 sheep (three replicates in each group) with different litter sizes, and H&E staining was performed for ovary structure observation. In the ovary, follicles were observed in different developmental states in both groups (Fig. 1a), which means that our subsequent analysis covers all cell types from various developmental states in the ovary during estrus.

**Figure 1.**
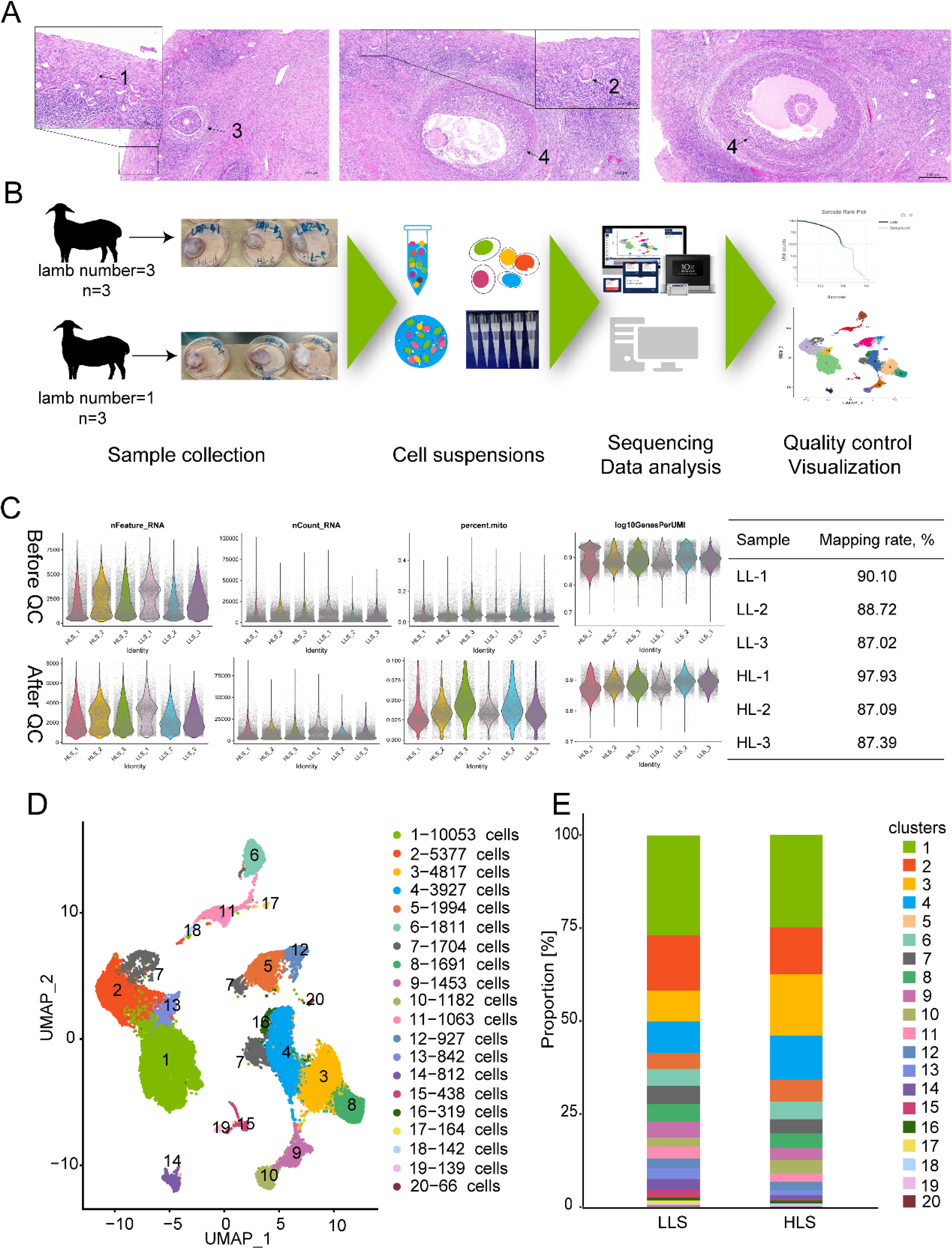
Single cell transcriptome sequencing of somatic cells in Hu sheep ovary. A, H&E staining of the ovary; 1, Primordial follicle; 2, 3, Growing follicle; 3, Antral follicle; 4, Graafian follicle. B, Procedure of ovary single-cell transcription sequencing. C, Quality control of single-cell transcriptome data. D, UMAP of ovary single-cell transcription sequencing data and clusters distribution in HLS and LLS groups.

The ovaries were digested for 10× genomic single cell RNA-seq (Fig. 1B). After critical cell filtration, 38921 cells were collected. The number of cells obtained from every sample ranged from 5796 to 7335, and the average number of UMI in each cell ranged from 6227 to 10384. The average number of genes in each cell ranged from 2160 to 3003, and the average proportion of mitochondrial UMI in each cell ranged from 0.0330 to 0.0480 (Fig. 1C). Mapping rate of every sample was higher than 85%, and the sc-RNA sequencing data of this study was reliable. Based on the sequencing data, a seurat-based workflow was used for cell clustering, and a total of 20 clusters (C) were identified by uniform manifold approximation and projection (UMAP) analysis. All clusters were present in HLS and LLS groups.

We characterized ovarian somatic cell types by existing cell markers provided by reference. The cells in C1, C2, C7-1, C13, and C14 were endothelial cells with high expression levels of marker genes, including *CDH5, CD34, VWF, FLI1,* and *MMRN1* [24]. GCs (C5, C7-2, C12) were recognized by the expression levels of *AMH*, *CDH2*, *FSHR,* and *FST* [25, 26]. According to the high expression levels of *PDGFRA, DCN,* and *TCF21*, cells in C3, C4, C7-3, C8, and C16 were identified as ovarian stromal cells [27]. Cells in C9 and C10 were annotated as perivascular cells based on the high expression of typical cell markers *RGS5, MCAM,* and *DES* [28]. The immune cells (C6, C11, C17, and C18) recognized by expression levels of *CD69, CD3G, PTPRC* and *CD53* [25]. The gene signatures of cells in C15, C19, and C20 were *DCDC2*, *MUC16*, *MPZ*, *CDH19*, *MZB1,* and *VPREB3.* The specific cell type related to these markers is rarely reported. Thus, these cells were identified as “unknown cells” (Fig. 2A). The immunofluorescence results also confirmed the expression and localization of different cell types (Fig. 2B, 2C). After cell type identification, five cell types of ovarian somatic from Hu-sheep were identified, and the cell number of endothelial cells, granulosa cells (GCs), stromal cells, perivascular cells, and immune cells were 17921, 3214, 11328, 2635 and 3180, respectively.

**Figure 2.**
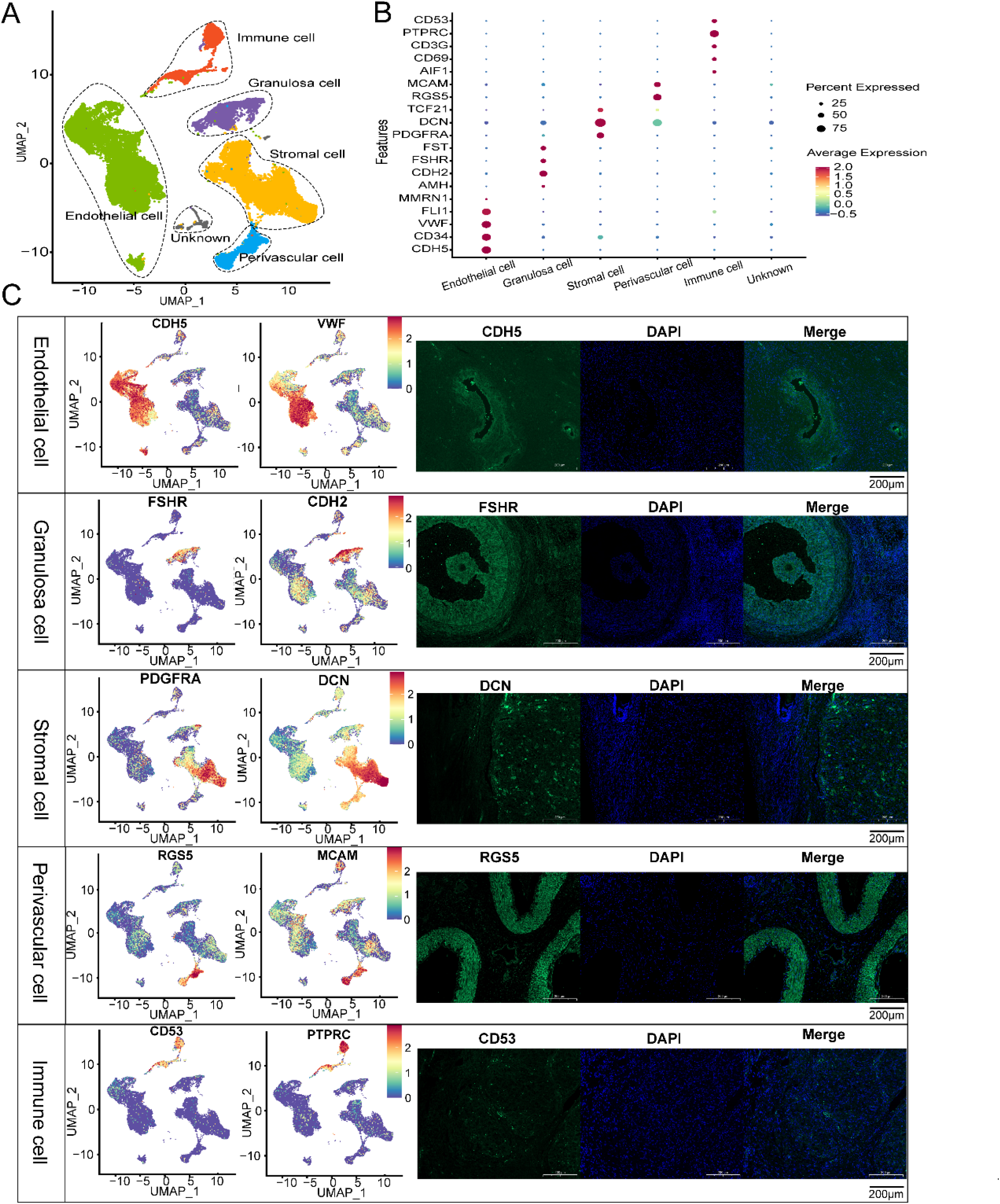
Identification of somatic cells in Hu sheep ovary. A, Identification result of five different cell types on UMAP. B, Dot plot of different cell marker gene expression levels. C, Representative marker genes’ feature plot and immunofluorescence of ovary somatic cell, green-gene, and blue-DAPI. Scale bar: 200 μm

### 3.3 Ovary somatic cell expression profiles during oestrum

We collected high-variable genes (Top 300) of each cell type for GO and KEGG enrichment analysis. The top 3 GO terms and KEGG pathways are shown in Fig. 3. GO and KEGG enrichment data uncovered the typical cell function and confirmed the cell type identification results. Numerous pathways related to metabolism (energy, amino acid, carbohydrate, and lipid) occurred in granulosa and stromal cells, which was indicated during oestrusm. GCs and stromal cells underwent a very energetic metabolism for follicle development and maturation. Some key pathways were enriched in certain cells during estrus. The AMP-activated protein kinase (AMPK) pathway is only enriched in GC. Forkhead box O (FoXo) signaling pathway is enriched in GCs and stromal. Protein kinase B (PI3K)-Akt signaling pathway enriched in endothelial cells and perivascular cells. The Mitogen-activated protein kinase (MAPK) signaling pathway was enriched for endothelial cells and immune cells.

**Figure 3.**
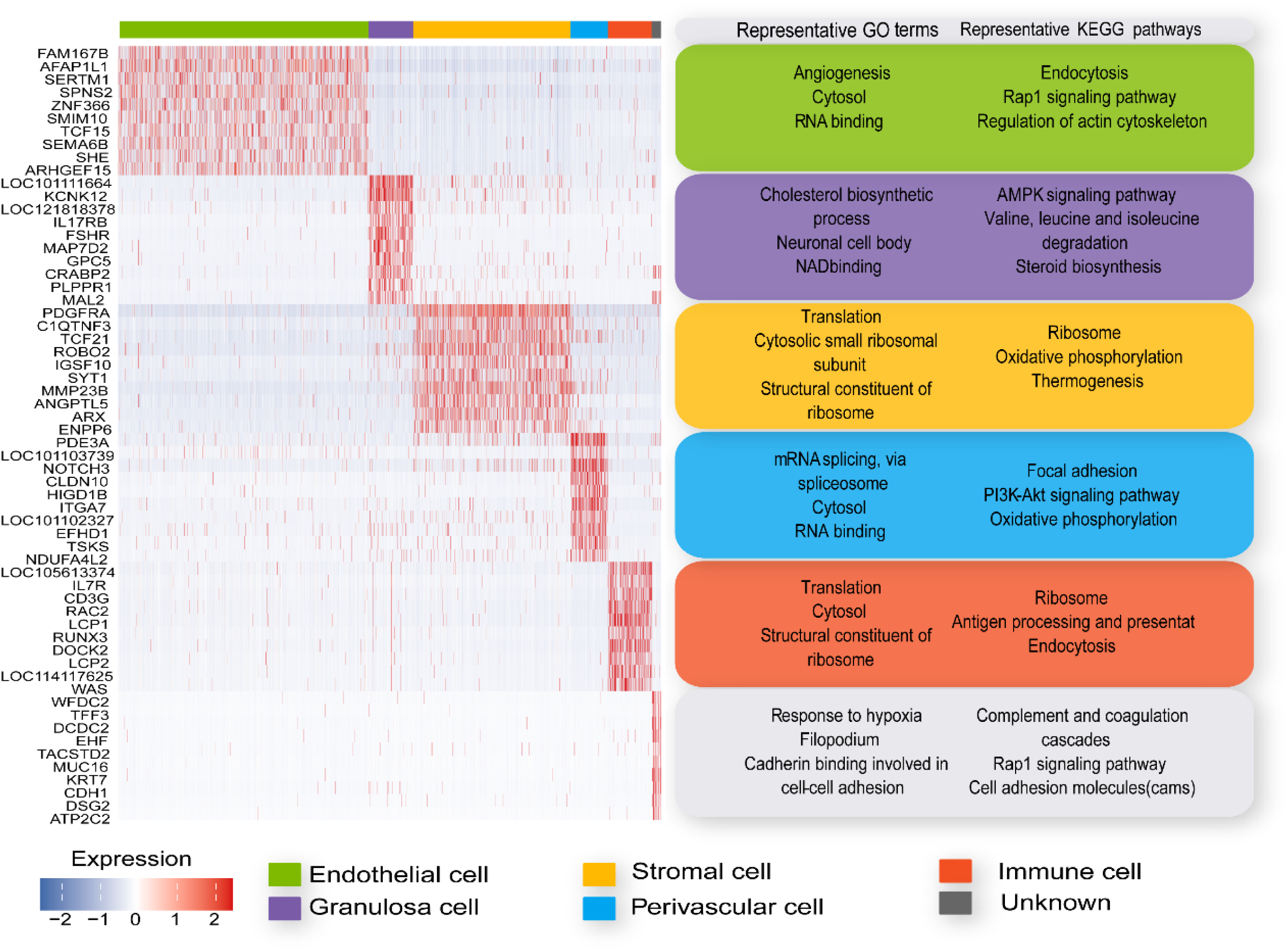
Ovary somatic cells marker genes heatmap and function enrichment.

### 3.4 Differences of ovary somatic cell expression profiles between HLS and LLS

The differences between the somatic cell expression profiles of single and multi-lamb Hu sheep were compared. In our study, the comparison of somatic cell expression profiles was based on the cell types. The proportion of cell types differed between HLS and LLS. The proportion of endothelial cells and immune cells in HLS was lower than in LLS, whereas GCs and stromal cells were higher than in LLS. The distribution of perivascular cells was consistent in both groups (Table. 3, Fig. 4A).

**Figure 4.**
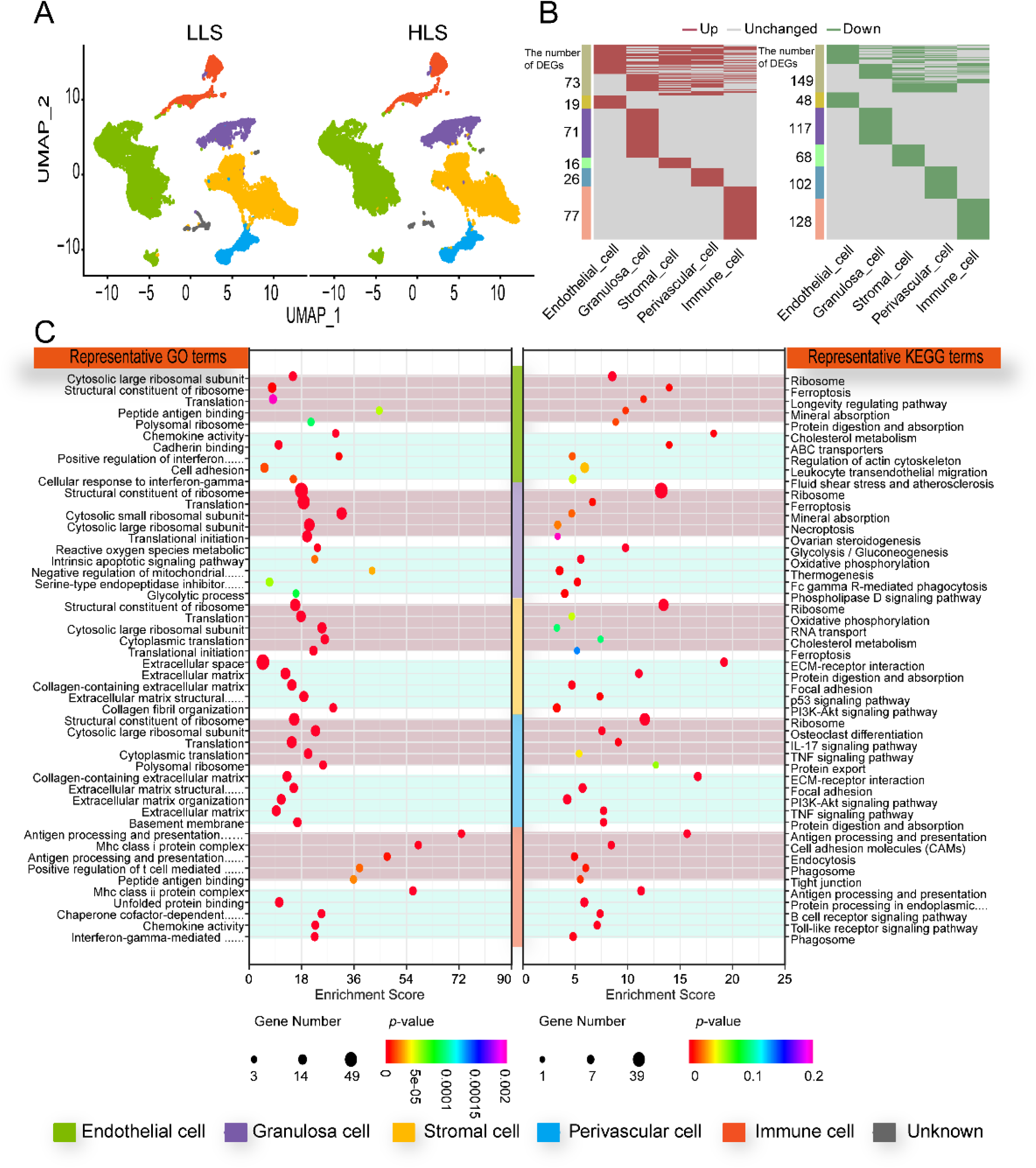
Comparison of differed cell type expression profiles between single and multi lamb sheep ovary. A, Somatic cell type difference in UMAP. B, Differentially expressed genes (DEGs) in a somatic cell of sheep ovary. C, Top 5 Gene Ontology (GO) and Kyoto Encyclopedia of Genes and Genomes (KEGG) enrichment, light red-up-regulated, light blue-down-regulated.

**Table 3.** Comparison of different cell type proportions between HLS and LLS.

| Cell type | LLS | HLS | *p*-value |
| --- | --- | --- | --- |
| Endothelial cell | 49.95±8.44 | 41.09±8.63 | 0.86 |
| Granulosa cell | 7.63±1.86 | 9.09±3.23 | 0.27 |
| Stromal cell | 23.89±4.52 | 34.74±3.55 | 0.61 |
| Perivascular cell | 6.59±0.48 | 6.88±0.67 | 0.54 |
| Immune cell | 9.1±1.74 | 7.53±3.02 | 0.29 |
| Unknown | 2.84±1.88 | 0.67±0.09 | 0.02 |

*p*-value<0.05 and Fold Change >1.2 were used as screening criteria. The numbers of up-regulated DEGs were 61, 115, 65, 74, and 105 in endothelial cell, GC, stromal cell, perivascular cell, and immune cell, respectively, while the down-regulated genes were 179, 168, 108, 181, and 179 (Fig. 4B). GO and KEGG enrichment analyses were conducted with DEGs. The enrichment results showed that the functions up-regulated in ovarian somatic cells were associated with ribosome-related functions compared to LLS. Upregulation of structural components of ribosomes, cytoplasmic large ribosomal subunits, translation, and other GO terms was observed in endothelial cells, GCs, stromal cells, and perivascular cells. The result of KEGG enrichment in ovarian somatic cells was closely involved in cellular functions and the enhanced ribosome pathway. The ovarian steroidogenesis pathway was up-regulated in GCs. Enhanced enrichment of the oxidative phosphorylation pathway was observed in stromal cells. Decreased functional enrichment of somatic cells was associated with extracellular structures and cell adhesion functions. For the GO term, cell adhesion enrichment was decreased in endothelial cells. Functional enrichment was decreased in stromal and perivascular cells regarding the extracellular matrix and its structural composition. In KEGG enrichment, a corresponding decrease in functional enrichment (ECM-receptor interaction) was observed in stromal and perivascular cells. The HLS group showed decreased nutrient metabolic functions in the KEGG enrichment results compared to the LLS group, such as decreased cholesterol metabolism enrichment in endothelial cells and decreased enrichment of protein digestion and absorption in stromal cells and perivascular cells. This difference was more pronounced in GCs, where the HLS group significantly downregulated enrichment of glycolysis/glucose production, oxidative phosphorylation, and thermogenesis. By comparing the expression profiles of ovarian somatic cells of Hu sheep with different lambing numbers, the number of differential genes and functional changes closely related to ovarian ovulation was greater in GCs than in other somatic cells. Thus, the difference in GCs in the different groups needs further analysis.

### 3.5 Granulosa cell subtype identification and developmental trajectory of sheep ovary

We reduced the dimension of the identified GCs into 11 sub-clusters (CL1-CL11) (Fig. 5A). CL1, CL3, CL5, and CL8 were recognized as early GC (eGC) through high expression of *WT1*, low expression of *VCAN* [29] [30], high expression of *TNNI3* [25] and *WNT6* [31]. Although we recognized these cells, the mapping condition was not ideal. So, we performed cell functions with GO and KEGG (Fig. S1). Based on the GO and KEGG enrichment results, we found the function related to signal transduction, response to peptide hormone and insulin, and the key pathways, such as Rap1 signaling pathway, FoxO signaling pathway, AMPK signaling pathway, Mammalian target of rapamycin (mTOR) signaling pathway and WNT signaling pathway enriched in CL1. Positive regulation of transcription by RNA polymerase II, nucleus, and DNA binding function was enriched in CL3. MAPK signaling pathway, steroid biosynthesis pathway, focal adhesion pathway, PI3K-Akt signaling pathway, and SMAD binding pathway were enriched in CL3. In CL5, functions related to the nucleus and RNA binding were enriched. The enriched pathways were the regulation of the actin cytoskeleton pathway, focal adhesion pathway, and relaxin signaling pathway. Function related to the nucleus, transcription corepressor activity, transcription factor binding and response to cAMP enriched in CL8, and pathways about PI3K-Akt signaling and MAPK signaling were enriched. Former studies have shown that in that WNT signal activation occurs exclusively at the primordial follicle stage [31]. Meanwhile, the FoXO signaling pathway, mTOR signaling pathway, MAPK signaling pathway, and PI3K-Akt signaling pathway have key functions in the activation of primordial follicles [32, 33]. Through the function enrichment results, we confirmed that these cell clusters belonged to eGC. Mural GC (mGC) (CL 2, CL9) was identified based on the expression levels of former reported cell markers *CITED2* [34], *FSHR* [35], *GJA4, IGFBP5,* and *CYP11A1* [36, 37]. CL4, CL6, and CL7 were cumulus GC (CC) since the high expression levels of marker genes *IHH, INH BB,* and *IGFBP2* [27], and CL10 and CL11 were recognized as atretic GC (aGC). The expression levels of *GJA1* and *CDH2* were lower in atretic follicles GC when compared with healthy follicles [38] (Fig. 5B, Fig. 5C).

**Figure 5.**
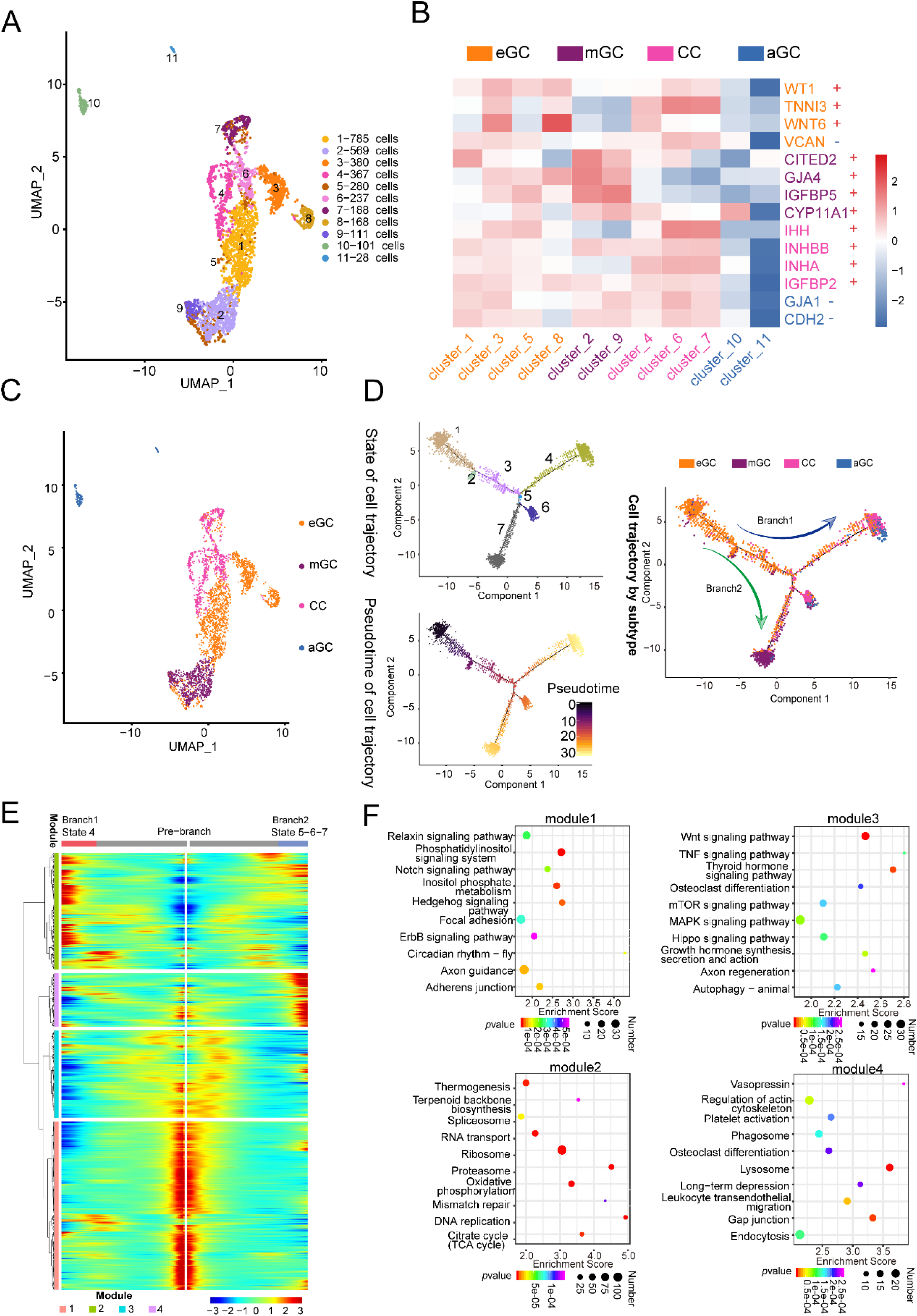
Granulosa cell subtype identification of sheep ovary. A, UMAP of reduced the dimension of granulosa cells. B, Granulosa cells sub-cluster marker gene heatmap. “-” indicates low expression of the corresponding gene. “+” indicates high expression of the corresponding gene. C, Granulosa cell subtype identification in UMAP. D, Granular cell pseudotime trajectory. E, Granulosa cell different developmental states heatmap. F, Top 10 KEGG enrichment of pseudotime heatmap genes.

We found a very extensive co-expression occurring in different GCs, which indicated that a lot of cell differentiation took place during oestrum. Although the ovaries were collected at a single time point, the special histology structure of the ovary could obtain follicles in different developmental states (Fig. 1A). Cell trajectory analysis explored the differentiation trajectory of GCs by Monocle. From trajectory analysis data, GC cells were divided into 7 states, both groups of GCs. All CLs were presented in trajectory. CL1 was presented almost in all states. CL2 was presented in states 6 and 7. CL3 was presented in states 1, 2, and 3. CL4 was presented in state 4. CL5 was presented in states 1 and 4. CL6 was presented in state 6. CL7 was presented in states 6 and 7. CL8 was presented in states 1 and 4. CL9 was presented in state 7, and CL 10 was presented in state 4 (Fig. S2). We identified the GCs’ sub-type and mapped the sub-type into the trajectory. The identified sub-type and the eGCs were located in the early state. The GCs were differentiated into two broad directions, including mGC and CC. Through trajectory analysis, in eGC, CL1 appeared in all states, and CL3 and CL8 appeared only in early development states, whereas CL5 intended to differentiate into CC (Fig. 5D). We performed heatmap analysis on cells in different developmental states and granular cells exhibited four patterns of gene expression levels. Genes in the pre-branch showed modle1 and 3 patterns, with high expression in the early developmental stages. Genes in branch1 and branch2 showed similar patterns of late high expression levels in the two different branches (Fig. 5E). KEGG and GO enrichment analyses of genes were conducted with different expression patterns, and the enrichment results of modle1 and modle3 were similar to those of eGC enriched in Adherens junction, Notch signaling pathway, WNT signaling pathway, Thyroid hormone signaling pathway, MAPK signaling pathway, Hippo signaling pathway, and mTOR signaling pathway. The growth hormone synthesis, secretion, and action pathway were enriched in modle1, indicating that cells were in a rapid growth period during this stage. In modle2, significant enrichment of functions was related to genetic material transfer, such as the ribosome, proteasome, DNA replication, RNA transport, and mismatch repair, as well as enhanced cell metabolism activities such as oxidative phosphorylation, thermogenesis, and citrate cycle, which suggests that cell metabolism is activated during the process of early GCs to cumulus cells development. In the enrichment results of modle4, functions related to the regulation of actin cytoskeleton, endocytosis, and phagosome were observed (Fig. 5F). These results justify the granulosa cell subtype identification strategy. The marker genes of each granulosa cell subtype were explored (Supplementary Material 2). The maker genes eGC, CC, and mGC were *WT1* and *CD34*, *AMH* and *INHA*, and *HTRA3*, respectively, for Hu sheep in our study.

To investigate key genes involved in the developmental timeline of granular cells, GeneSwitches analysis was conducted. Along the timeline of the branch1 developmental process, we observed gene closure throughout the entire timeline, leading to the development of early GCs into cumulus cells. In the later stage, more transcription factors, such as *JUNB* and *FNDC3B*, and a key gene, such as *FOSB*, participated in the closure process. At 23.4 h, *LOC101112291* expression was closed. These genes decreased expression levels during the final transformation into cumulus cells (Fig. 6A, Fig. 6C). Along the timeline of branch2, the expression levels of genes increased in the early stage, with transcription factors (*UBA52* and *PPIB*). Some patent factors are related to energy metabolisms, such as *ACTG1, LDHB, ATP5MG, ATP5MC3*, and *ATP5F1E*, opening up in expression. In the late stage of the timeline, we observed the opening of *JPH1* expression. These genes were expressed higher when early GCs were transformed into mural GCs (Fig. 6A, Fig. 6C).

**Figure 6.**
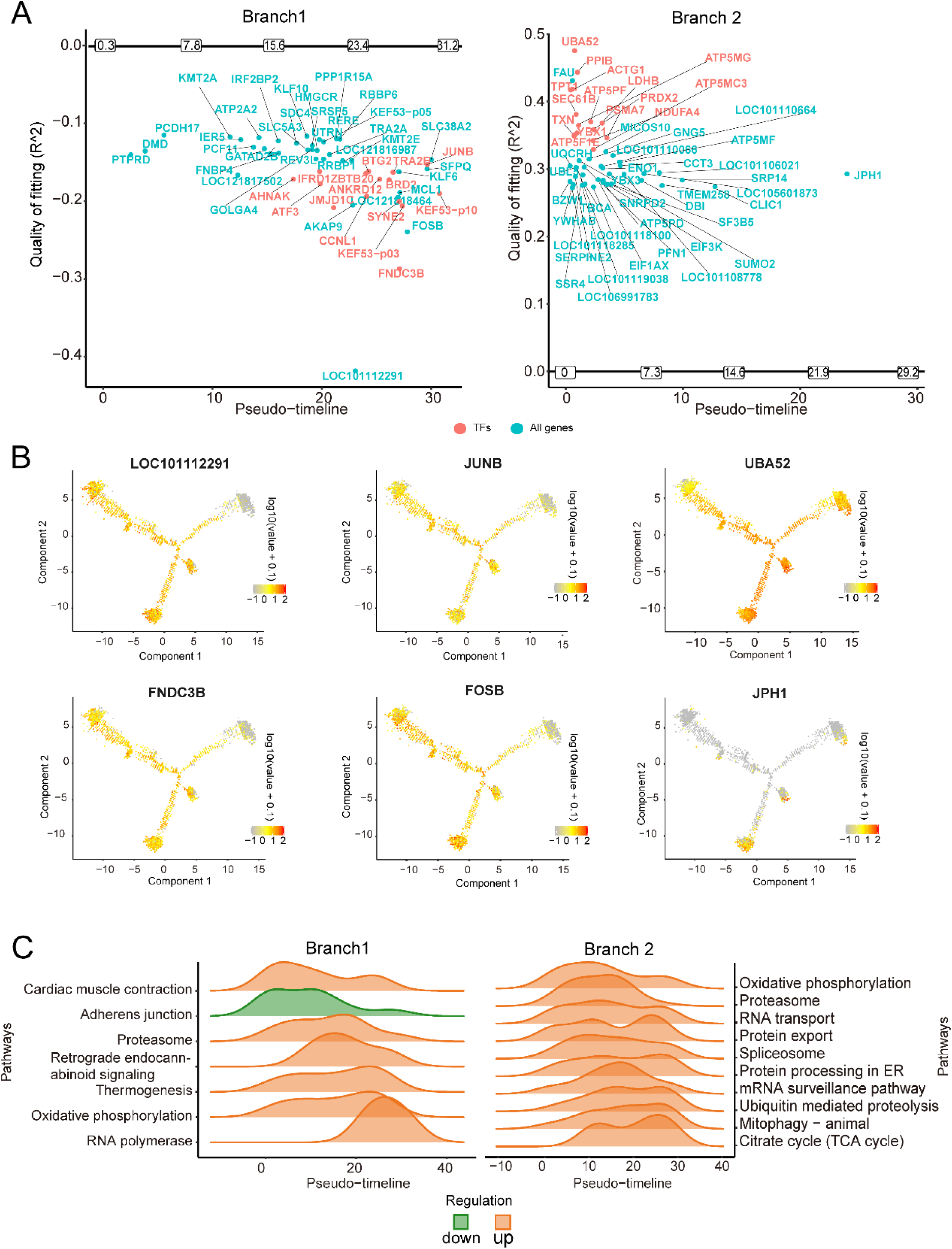
Granular cell developmental trajectory with pseudotime. A, Key genes involved in the developmental process of granular cells. The horizontal axis is pseudotime, and the vertical axis is the goodness-of-fit R^2. The genes turned on with the pseudotime are shown above the horizontal axis, and the genes turned off are shown below the vertical axis. Genes that satisfy the following conditions have been selected for mapping: 1. The percentage of zero-expressing cells is below 90%, 2. The top 50 plots with the highest goodness-of-fit. B, feature plot of representative gene expression. C, Top 10 KEGG enrichment of genes involved in granular cells developmental process. Up, function enrichment turned on; down, function enrichment closure.

We performed KEGG pathway enrichment analysis on key genes in different branches. In the process of granulosa cell developing to CC (branch1), only adherens junction pathway closure. The other pathways were turned on. Adherens junction pathway was enriched at lower levels in the early stages of development and decreased with the timeline, while proteasome, thermogenesis, and oxidative phosphorylation increased with the timeline and decreased rapidly before the endpoint. RNA polymerase function only increased in the late stages of development. In the process of granulosa cell to mGC (branch2) development, the enrichment of pathways was turned on, focusing on cell energy metabolism (oxidative phosphorylation, TCA cycle) and related functions of cell protein synthesis (RNA transport, protein export, and protein processing in the endoplasmic reticulum) (Fig. 6C).

### 3.6 Differences of ovary granulosa cell expression profiles between HLS and LLS

By comparing the proportions of different cell subtypes, the proportion of atretic follicular granulosa cells (aGCs) was significantly higher in the LLS group than in HLS (Fig. 7A). In the LLS group, the physiological status of GCs was altered, resulting in an increase in follicles with a propensity for atresia. Changes in cell function based on cell subtypes were analyzed using KEGG enrichment; pathways associated with apoptosis and necroptosis were inhibited, while pathways associated with cell survival were up-regulated. This change was cell subtype based. In mGCs, the enrichment of the necroptosis pathway was elevated. The genes enriched in the pathway were *FTH1, FTL,* and *H2AZ1*. *FTH1* and *FTL* in the necroptosis pathway had a negative feedback regulatory effect in the ferroptosis pathway. In the current study, *FTH1* and *FTL* highly expressed in group HLS reduced necroptosis by reducing the release of reactive oxygen species (ROS) after decreased lysosome membrane permeabilization. The expression levels of ferroptosis-resisted genes *FTH1* and *FTL* were up-regulated in HLS. Then, we retrieved the location of *FTH1* and *FTL* on the ferroptosis pathway and analyzed its upstream and downstream genes. The downstream expression levels of key genes *MAP1LC3A, ATG5, ATG7* (Fig. S3), and their complex gene *NCOA4* (Fig. S3) were downregulated in the HLS group, thus reducing ferroptosis via inhibiting the Fenton response. On the other hand, the enrichment of the FoxO signaling pathway was downregulated in HLS and reduced apoptosis by decreasing the expression of *IRS2, EP300, BCL2L11*, and *SGK1* (Fig. S3). In CCs, the enrichment of ECM-receptor interaction in the multi-lamb group was up-regulated by increasing the expression levels of *COL4A4* (Fig. 7D), whereas the enrichment of the cAMP signaling pathway, oxytocin signaling pathway, tight junction, and thyroid hormone signaling pathway was down-regulated in the multi-lamb group, with decreasing the expression levels of *CALM1* (Fig. 7D)*, PLD1, OXT, PLN, ATP2B1, F2R, ACTG1,* and *MYL6* (Fig. S3). In eGC, differences between the two groups were reflected in the down-regulation of the expression of transcription factor complex AP-1 composed of *FOS, FOSB,* and *JUN* (Fig. 7B, Fig. 7C).

**Figure 7.**
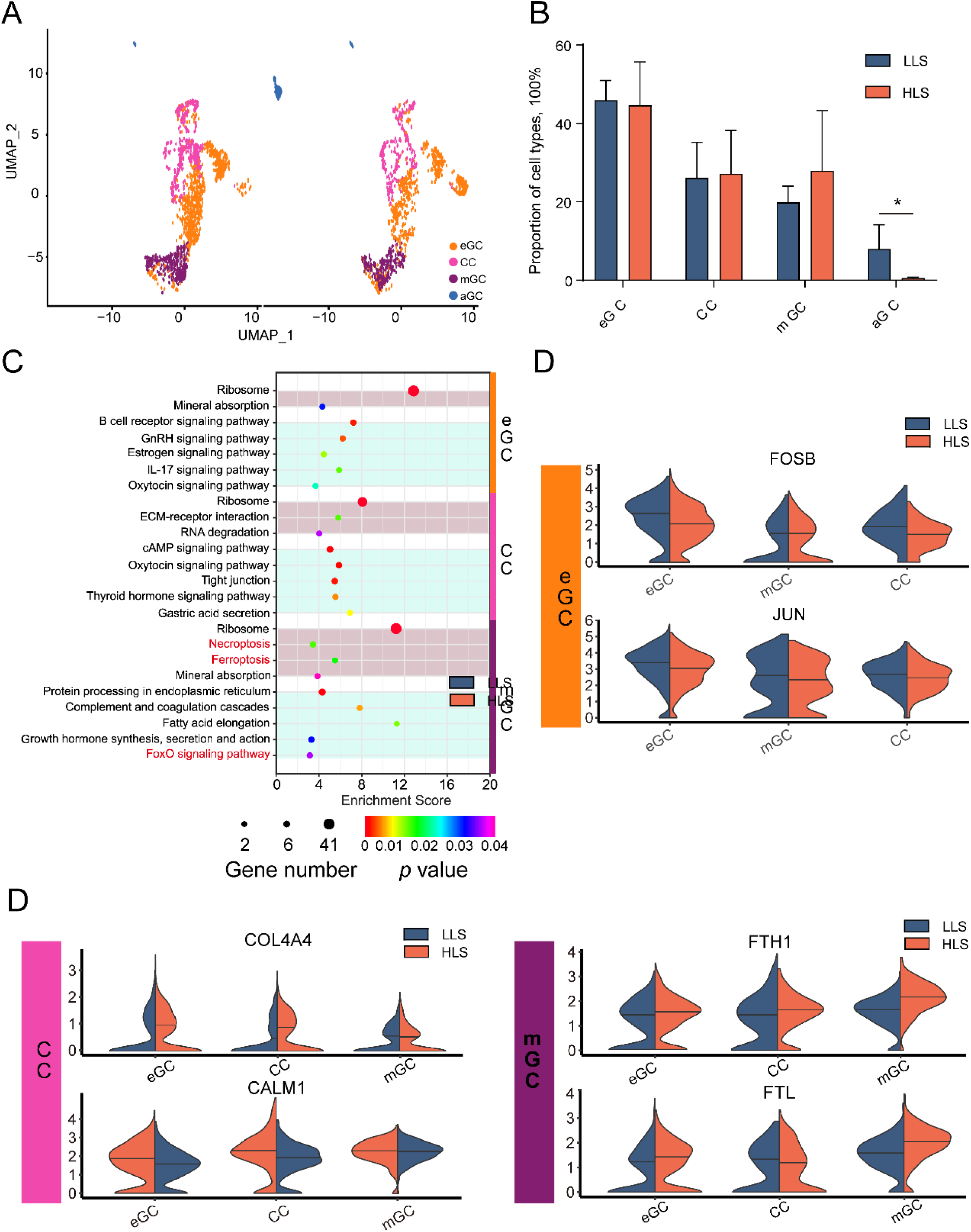
Comparison of different granulosa cell sub-type expression profiles between single/mult lamb sheep ovary. A, Granulosa cell sub-type difference in UMAP. B, Comparison of different granulosa cell sub-types proportion between HLS and LLS of cell Differentially expressed genes (DEGs) in a somatic cell of sheep ovary. C, KEGG enrichment of DEGs in ovarian granulosa cell subtypes of sheep with different lambing numbers, light red-up-regulated, light blue-down-regulated. D, Comparison of representative genes of ovarian granulosa cell subtypes in sheep with different lambing numbers.

## 4. Discussion

In sheep’s ovary, every estrus cycle, there are typically 3 or 4 waves of follicle development during the inter-ovulatory interval [39], and 1∼3 mature follicles ovulate [40]. Estrus is a special time window. In peripheral blood, LH peaks can be observed; E2 decreases rapidly from maximal values; progesterone is at its lowest level, and sheep usually ovulate about 20 h after the onset of estrus [41]. Thus, understanding the transcriptional profile of ovarian somatic cells during estrus is essential to investigate the mechanism of ovulation. In the present study, we investigated the differences in expression profiles of single/multi-lamb Hu sheep ovary somatic cells by single cell RNA seq, providing insight into the mechanisms underlying the differences in ovulation numbers. A total of five types of somatic cells were identified, and corresponding expression profiles were mapped in the ovaries of Hu sheep. We performed subtype identification of GCs was performed. Key genes involved in different subtype transitions were analyzed. The differences in cells’ expression profiles were compared to clarify the key factors regulating different litter sizes. These findings give a theoretical basis for breeding high-fertility sheep and provide new targets for molecular genetics-based selection.

### 4.1 Identification of ovary somatic cells

Various cells in the ovary act in synergy to enable ovarian function, whereas existing research has not paid much attention to the function of these somatic cells, except GCs. The ovarian stroma comprises mostly incompletely characterized stromal cells (e.g., fibroblast-like, spindle-shaped, and stromal cells) [42]. In recent years, the role of stromal cells in the ovary has been revisited, and studies have identified estrogen receptors α and β in the cytoplasm and nucleus of bovine stromal cells, unlike fibroblasts. These cells are oval cells with lipid droplets and vacuoles [43]. Progesterone receptor α has been identified in stromal cells of pregnant and postpartum rabbit ovaries [44]. In the present study, a large amount of energy metabolism occurred in the ovaries during estrus for supporting ovulation. There is an extensive blood supply to the ovary, which is involved in forming dominant follicles, and the endothelium can participate extensively in the angiogenic process. The importance of combined transplantation of ovarian endothelial cells with stromal cells when performing follicular transplantation in individuals with premature ovarian failure was demonstrated to ensure the formation of a well-vascularized and well-structured ovarian-like stroma [45]. A previous study proposed that perivascular cells were multipotent progenitors that contribute to granulosa, thecal, and pericyte cell lineages in the ovary, which supports folliculogenesis [46]. The main functions of immune cells in the ovary are defense, remodeling of ovarian structure, signaling [47], and ovarian aging [48]. In the present study, marker genes of ovary somatic cells proposed were confirmed in former studies [28]and also available as stroma cell gene signatures of Hu sheep.

Among the ovarian somatic cells, the most well-studied are the GCs. The GC is a somatic cell surrounding the oocyte co-located with the oocyte in the same follicular microenvironment. Its function is limited to the secretion of gonadotropins to stimulate ovulation and includes follicular development. GCs secrete factors, including gonadal steroids, growth factors, and cytokines are critical for GC survival and follicular growth [49, 50]. In contrast, identifying the GC subtype remains controversial, especially for sheep. In the human ovary, the expression pattern of early-stage GCs is *WT1*^high^/*EGR4*^high^/*VCAN*^low^/*FST*^low^. The expression pattern of cumulus GC is (*VCAN*^high^/*FST*^high^/*IGFBP2*^high^/*HTRA1*^high^/ *INHBB*^high^/*IHH*^high^), and the expression pattern of mural GC is *WT1*^low^/*EGR4*^low^/ *KRT18*^high^/*CITED2*^high^/*LIHP*^high^/*AKIRIN1*^high^ [27]. In domestic animals, the GCs of goats were identified based on developmental trajectory. *ASIP* and *ASPN* were highly expressed in early GCs, *INHA, INHBA, MFGE8,* and *HSD17B1* were highly expressed in GCs during the growth phase, and *IGFBP2, IGFBP5,* and *CYP11A1* were highly expressed during the growth phase of GCs [26]. However, the study did not give subtype classification markers based on cell function. The present study defined GC subtypes by combining existing marker genes and functional analysis of different sub-clusters. Using pseudotime analysis, the reliability of GC subtype identification was verified. It has been found that *WT1* and *CD34* are marker genes for eGCs. *AMH* and *INHA* are marker genes for CCs, and *HTRA3* is a marker gene for mGCs. These marker genes were applicable for identifying sheep GC subtypes.

Five somatic cell lineages were identified in sheep ovaries based on their gene expression signatures. GCs were further characterized into three subtypes, marker genes for each cell type are only expressed in specific “regions” in the UMAP figure and immunofluorescence profiling, which were consistent with the anatomy of the ovary [51]. These results illustrated the reliability of the single-cell sequencing data from this study. However, no luteal cells were detected in our dataset, which is consistent with our previous study [24], which implies the degradation of luteal cells during the samples collection period (estrus) or the luteal cells are difficult to collect because the cell extraction methods used in existing studies.

### 4.2 The transition of different GC subtypes

CC and mGC interact with oocytes differently in the follicle. The CC carries out bidirectional information transfer with the oocyte through gap junctions, contributing to oocyte maturation, fertilization, and early embryonic development [52]. In contrast, the mGC has multiple receptors on its surface that can secrete various hormones and cytokines that regulate follicular growth and maturation in an autocrine and paracrine manner [53]. In the present study, key genes were observed which are involved in the transition of different GC subtypes using GeneSwitches. The closure of *LOC101112291* led to the differentiation of eGCs into CCs, while the opening of *JPH1* expression led to the differentiation of eGCs into mGCs. A previous study investigating the molecular mechanism of lambing in Hanper sheep using ovarian tissue has revealed that *LOC101112291 (XIST)* regulates lambing number through the methylation process [8]. On the other hand, the protein expressed by the *JPH1* gene, Junctophilins (JPHs), is a family of structural proteins that connect the plasma membrane with intracellular organelles such as the endoplasmic/sarcoplasmic reticulum (ER/SR). The anchoring of these membrane structures leads to highly organized subcellular connections, playing an important role in signal transduction in all excitable cell types [54]. Our study found that the expression levels of these genes were turned off. Therefore, *LOC101112291* and *JPH1* genes may potentially regulate the direction of differentiation of early GCs.

### 4.3 Differences in transcriptional profiles of GCs in Hu sheep with different lambing numbers

In the modern sheep production system, the reproductive performance of female animals determines the economic profitability of farming, and how to increase the number of lambs has always been the hottest spot and key in sheep genetic breeding and reproduction research. Based on former studies, GC is vital in follicle development [49, 50, 55]. Our data revealed the number of differential genes and the key functional differences in single/multi-lamb distributed in GCs, so we paid attention to these cell clusters. Li et al. [26] studied the gene expression of GCs at different stages in 2 populations of Jining gray goats, and they found differences in the enrichment of GO terms of GCs at different periods in different lambing numbers groups. The previous study showed the differences in the expression profiles of GCs at different lambing numbers from functional analysis. In this study, the definition of the subtypes of Hu sheep GCs enabled us to discover differences in the functions of GCs in the two groups. Follicular atresia was increased in the LLS group, which was mainly caused by ferroptosis of GCs. Healthy growing follicles have a granular layer that is aligned with the follicular basement membrane, and no apoptotic cells are present. In the early stages of follicular atresia, apoptotic GCs gradually appear and increase in number. In progressive atretic follicles, most GCs undergo apoptosis leading to severe disruption of the granular layer and clearance of the follicle. Apoptosis is initiated in the GCs on the inner surface of the granular layer, while the oocytes, as well as the inner and outer layers of the membrane, are not affected by apoptosis in the early stages of atresia [56], suggesting that GC apoptosis plays an initiating role in follicular atresia [57, 58]. Ferroptosis is a form of cell death caused by iron-dependent lipid peroxidation and ROS accumulation characterized by the reduction or loss of mitochondrial cristae and rupture of the outer mitochondrial and mitochondrial membranes condensation [59]. Zhang et al. [60] found that transferrin (TF) expression was significantly reduced, and *PCBP* expression was significantly increased in porcine early atretic follicles, suggesting that iron accumulation began to occur early in follicular atresia and ferroptosis had an essential regulatory role in follicular atresia. Another study on female infertility found that induced iron overload in GCs led to ferroptosis and suppressed oocyte maturation by releasing exosomes from GCs, suggesting that ferroptosis of GCs is detrimental to oocyte development [61]. This study found that the GCs of mult-lamb sheep suppressed ferroptosis by increasing the expression levels of anti-ferroptosis genes *FTH1* and *FTL*, which promotes oocyte maturation and prevents follicular atresia, contributing to the mult-lamb trait.

## 5. Conclusion

In our study, we identified differences in the expression profiles of ovarian somatic cells between high-litter and low-litter size Hu sheep. These differences were mainly attributed to granulosa cells. The expression condition of *JPH1* and *LOC101112291* emerged as a significant indicator for determining the evolutionary directions of granulosa cells. Additionally, *FTH1* and *FTL* were identified as potential genes that regulate litter size. This study provides new insights into the molecular mechanisms underlying the high reproductive rate of Hu sheep.

### Declarations

#### Ethics approval and consent to participate

The protocol for this study was approved by the Animal Care and Use Committee of Northwest A&F University, China (Approval No. DK2021113).

The methods for this study were carried out in accordance with relevant guidelines and regulations of Northwest A&F University, China (Approval No. DK2021113).

All methods are reported in accordance with ARRIVE guidelines (https://arriveguidelines.org) for the reporting of animal experiments.

#### Consent for publication

All authors have read and approved the manuscript.

#### Author’s contributions

Ting Ge wrote the main manuscript text and performed single-cell RNA sequencing and corresponding data collection, analysis, and visualization. Ting Ge and Yifan Wen performed blood biochemical and hormone levels tests, hematoxylin and eosin staining, and immunofluorescence staining. Yifan Wen, Xiao Yu Huang, and Bo Li conducted data analysis and edited the manuscript. Ting Ge, Shaohua Jiang, Xiao Yu Huang, and Bo Li collected the samples. En Ping Zhang supervised the entire experiment.

## Acknowledgements

In this research, we acknowledge all members of the Innovative research team of sheep and goats of Northwest Agriculture & Forestry University.

## Author’s information

Affiliations: College of Animal Science and Technology, Northwest A&F University, Xianyang 712100 Shaanxi, China

